# Persistent articular infection and host reactive response contribute to *Brucella*-induced spondyloarthritis in SKG mice

**DOI:** 10.1101/2025.02.18.638825

**Authors:** Jerome S. Harms, Michael Lasarev, Thomas Warner, Sergio Costa Oliveira, Judith A. Smith

**Author notes:** Address correspondence to Judy Smith,. deceased (Thomas Warner).

## Abstract

Brucellosis, one of the most prevalent zoonotic diseases worldwide, often results in osteoarticular complications including large joint and axial arthritis mimicking spondyloarthritis. To model this chronic manifestation, we infected autoimmunity-prone SKG mice containing a mutation in the T-cell adaptor ZAP-70 with *Brucella* species. *B. melitensis* infection resulted in a fully penetrant, readily scoreable disease involving large joint wrist and foot arthritis, peri-ocular inflammation, and less frequent scaly paw rash. Infection with *B. abortus* resulted in delayed arthritis onset, and *B. neotomae* revealed sex differences, with more severe disease and a dose response in females. Heat-killed *Brucella* did not induce arthritis, evincing a requirement for viable infection. Across species, splenic CFU correlated well with final clinical score at 12 weeks (ρ=0.79 and p<0.001). *In vivo* imaging using luminescent *B. neotomae* revealed rapid colonization of the paws by one-week post-infection, more than a month prior to arthritis onset. Paw luminescence levels decreased after 2 weeks and then remained relatively static, even as clinical scores increased. Thus, the degree of arthritis did not strictly correlate with degree of paw infection but suggested an additional reactive component. Further, in examining a *Brucella* Δ*tcpB* mutant lacking a Type IV secretion system-dependent mediator, mice displayed an intermediate phenotype without significant differences in splenic CFU. Together these data suggest *Brucella* induced spondyloarthritis reflects both persistent colonization as well as excess host reactivity. Moreover, the sensitivity of the SKG model to different species and mutants will provide new opportunities for dissecting correlates of *Brucella* virulence and host immunity.

**Importance:** Brucellosis, a bacterial infection acquired from herd animals, remains one of the most common zoonotic diseases worldwide. Chronic infection often results in spondyloarthritis-like complications. Investigation into pathogenesis has been limited by the lack of overt disease in standard lab mice. We addressed this issue using spondyloarthritis-susceptible SKG mice. Upon infection with *B. melitensis*, SKG mice develop robust, fully penetrant large joint arthritis. Arthritis development required viable bacteria. Moreover, studies of colonization, gene expression and anatomic distribution using bioluminescent bacteria revealed active persistent infection in the mouse paws. However, peak paw infection occurred much earlier than arthritis onset, suggesting an added immune reactive component. Disease onset, severity and manifestations varied upon infection with different *Brucella* species and mutants. Together these results suggest this new model will be very useful to the scientific community for determining correlates of bacterial virulence leading to clinical disease.

## Introduction

Brucellosis, caused by the facultative intracellular bacteria *Brucella* species, remains one of the most prevalent zoonosis worldwide, afflicting people in S. America, Mexico, Europe (Mediterranean basin), the Middle East, Africa, and Asia^1,2^. Initially, infection manifests with undulant fever, myalgias and arthralgias. However, chronic brucellosis may affect many host organs, leading to arthritis, orchitis, liver damage, endocarditis, and encephalomyelitis^3,4^. Osteoarticular involvement (sacroiliitis, peripheral arthritis, spondylitis, osteomyelitis, tenosynovitis) is the most frequently reported complication, affecting up to 20-87% of subjects^5,6^. Indeed, differentiation of chronic Brucellosis from rheumatologic spondyloarthritis can be challenging in endemic areas. *B. melitensis*, which infects sheep or goats, is the primary human pathogen, although human infections with *B. abortus* (cows) predominates in certain locations, and *B. suis* (pigs) infections also occur^2,7^. Of these species, *B. melitensis* is considered the most virulent species in humans, and associates with the more severe manifestations^2,8^. Brucellosis is challenging to treat, requiring prolonged courses of multiple antibiotics and relapses occur in 5-10% of cases^3,4^.

The study of *Brucella* osteoarticular disease has been hampered by limitations in current mouse models. BALB/c and C57BL/6 mice, the most commonly used *Brucella* models, develop a transient spread of *Brucella* to their tails and paws 7-10d post infection^9^. However, this colonization is mostly subclinical and requires high doses of luminescent *Brucella* and biophotonic imaging to detect^9–11^. Chronically infected BALB/c females have been reported to develop peripheral reactive arthritis 26 weeks post-infection, but with extremely low penetrance^11^. IFN-γ-/- mice develop arthritis after intra-peritoneal injection, but IFN-γ deficiency severely compromises *Brucella* containment^12^. Similarly, the NOD-SCID model develops destructive tail osteomyelitis in response to the vaccine strain S19, but completely lacks an adaptive immune system^13^. Foot pad and joint injections of bacteria do not capture the systemic and chronic facets of disease^14^.

To address these limitations, we turned to a murine model frequently used to investigate rheumatologic spondyloarthritis, the “SKG” mouse. Named for their discoverer (Shimon Sakaguchi), SKG mice bear a spontaneous mutation in the T cell receptor signaling adaptor ZAP-70 (W163C) that generates an autoimmunity-prone T cell repertoire. Mice in specific pathogen free conditions do not develop disease without a trigger. Injection of the fungal cell wall component curdlan (beta 1,3-glucan) results in enthesitis, peripheral and axial arthritis, vertebral inflammation, ileitis, and uveitis^15^. The curdlan-injected mice develop clinically obvious disease in a few weeks, with greater severity and rapidity in females. Intriguingly, this model is also dependent upon IL-23 and the presence of fecal microbiota^16,17^. SKG mice have also proven useful in modeling TNF-dependent *Chlamydia*-induced reactive arthritis^18^. Finally, the SKG mice are on the BALB/c background, which is more susceptible to *Brucella* than other strains^19^.

In this study, we investigated the clinical responses of SKG mice to different species and attenuated mutants of *Brucella*. Additionally, we sought to determine if the arthritis represented a purely “reactive” response to the initial infectious stimulus or persistent infection. Together our results suggest the severity of disease development reflects contributions both from persistent systemic infection as well as excess host reactivity. This new *Brucella* SKG model should prove highly useful for probing *in vivo* correlates of bacterial virulence and host immune responses to infection that result in spondyloarthritis manifestations.

## Results

### *B. melitensis* induces spondyloarthritis-like clinical manifestations in SKG mice

Mice were infected intraperitoneally with *B. melitensis*, as this route results in systemic dispersal of the bacteria mimicking human disease, including seeding of the spleen, liver, placenta, testes and the skeleton^19^. No effects of infection were observed until after 3 weeks, but by 4 weeks (“chronic” phase for mouse infection), 100% of the mice began developing arthritis in large weight-bearing joints (wrists and ankles/feet) with symmetric involvement (**Figure 1A and 1B**). The hind paws exhibited mid-foot swelling reminiscent of tarsitis in human spondyloarthritis subjects. Small joints (paw digits) were spared. The second most prevalent manifestation was bilateral peri-ocular inflammation with an initial waxing and waning course, occurring after 5-6 weeks. Around 8 weeks, a few animals developed scaly rash on their paws. Tail swelling was rare. In the first experiment a couple of control females developed mild late disease, but none in subsequent experiments. Infected animals also stopped gaining weight (**Figure 1C**). Based on the 2 most common manifestations (eye disease and arthritis), mice were scored for clinical disease severity (Methods, **Figure 1D**). Paw widths were also measured using calipers and revealed separation between infected animals and controls for end-experiment paw widths, and paw width changes over time (**Figure 1E**). Clinical scores were less variable than the caliper measurements, effectively compressing variation, and the arthritis scoring correlated well with changes in caliper measured paw widths (ρ∼0.76, p<0.001, **Supplemental Figure 1**) and end paw widths (ρ∼0.69, p<0.001, not shown). There were no differences between males and females for clinical score or changes in paw width in response to infection with *B. melitensis*. Histologic sections of paws at 9.5 weeks post-infection showed extensive bony destruction, areas of florid fibrosis, foci of neutrophils resembling chronic abscesses, lymphocytic foci, as well as lymphocytic and macrophage infiltrate extending into the surrounding tendons and muscle (**Figure 1F**). Although some mice developed transient diarrhea, gut histology by this point was normal. Strikingly, using micro-CT to visualize the skeleton, 12-week paws exhibited both extensively moth-eaten bones adjacent to the wrist as well as exuberant new bone formation. New bone formation is a hallmark of human rheumatologic spondyloarthritis. Interestingly, the phalangeal and carpal bones were relatively spared (**Figure 1G**).

**Figure 1.**
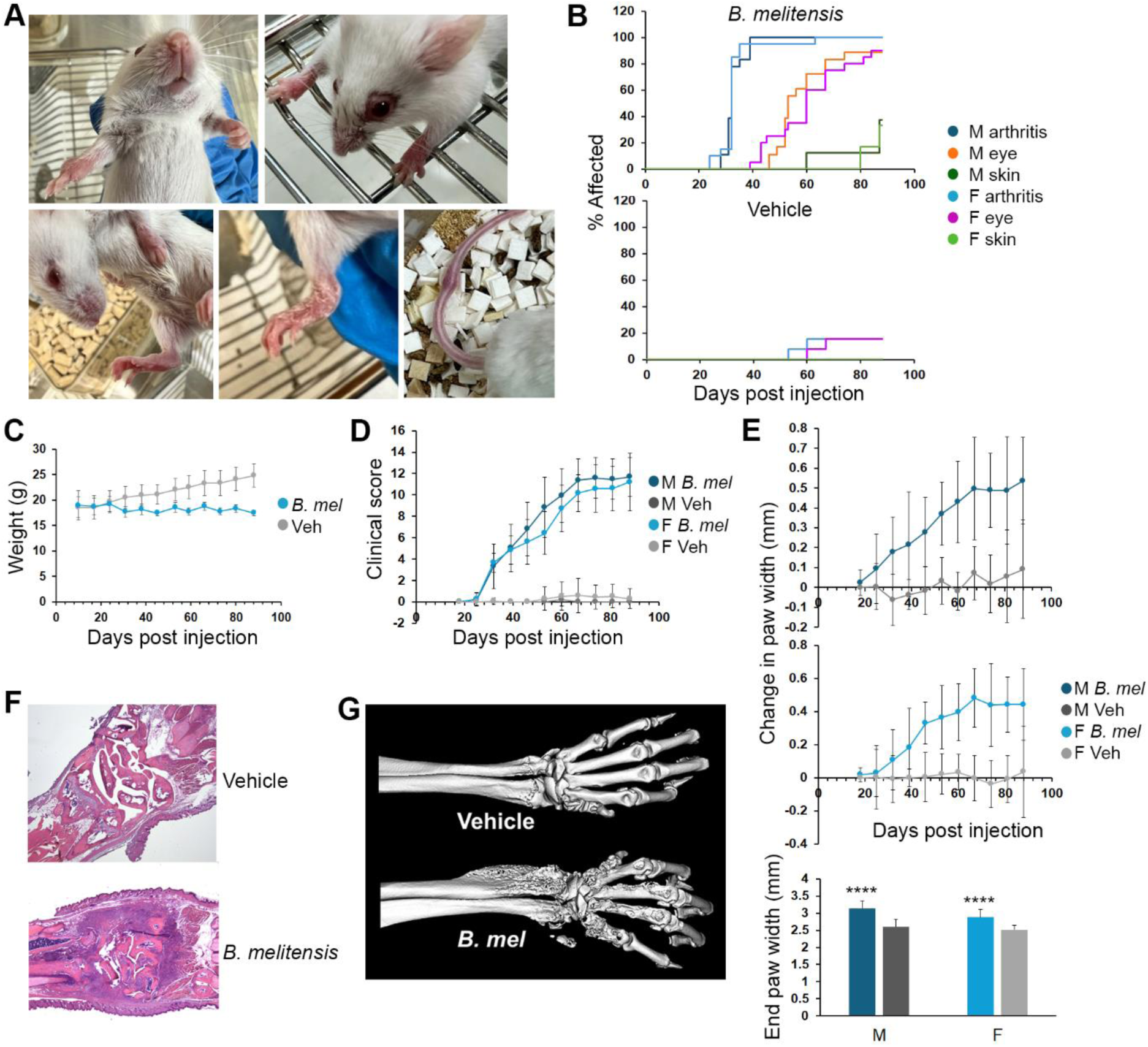
Clinical manifestations of *B. melitensis* infection in SKG mice. Mice were injected with PBS (vehicle control) or 2×10^6^ *B. melitensis* intraperitoneally. Results are aggregates from at least 3 experiments and include 18 infected males (M), 12 control males, 19 infected females (F) and 13 control females. A) Photographs of *B. melitensi*s infected mice showing (top left then clockwise): front paw arthritis, peri-ocular inflammation, tail swelling, scaly rash, mid-foot swelling. B) Percent of mice developing arthritis, peri-ocular inflammation and rash over time in male and female mice. Eye and skin disease occurred later than arthritis (p<0.001) C) Weights (grams) from infected or control female mice (N=6 and 4 mice, respectively, p=0.028 for weight loss in infected animals vs p<0.001 for weight gain in controls, p<0.001 for comparison). D) Mice were scored weekly for arthritis and eye inflammation as described in the methods. After day 25, p<0.001 for *B. melitensis* infected vs. control mice at each time point and trajectory over time. E) Front and hind paw widths were measured in mm with calipers and the four paws averaged for each mouse. The female infected group includes 14 mice. Changes in paw widths are vs. day 14-18 of the experiment. After day 25, p<0.001 comparing average paw widths for infected vs control mice at each time point, and rate of change over time. End paw widths are the average per mouse on Day 84-88 of the experiment. P<0.001 vs control, with no difference between sexes. F) Representative histologic sections from PBS (top) or *B. melitensis* (bottom) infected mice at 9.5 weeks post-infection. G) Micro CT images from PBS (top) or *B. melitensis* (bottom) infected mice at 12 weeks post-infection. ****p<0.001.

### Clinical manifestations following infection with other *Brucella* species

In comparing infection between *B. melitensis* 16M and *B. abortus* 2308, non-significant trends were noted in median onset of eye inflammation and paw rash (earlier onset, p=0.11 and p=0.06 respectively), but arthritis occurred about a week later with *B. abortus* (p<0.001, **Figure 2A**). Changes in paw width diameters following *B. abortus* were significantly different vs. control PBS-injected animals (p<0.001 for trends over time, **Figure 2B**). End paw widths did not differ between *B. abortus* and *B. melitensis* but were greater than controls. Although *B. abortus* induced milder disease in some mice (see clinical score box-whisker plots for male mice infected with *B melitensis* or *B abortus*, **Figure 2C**), the variation in clinical scores was not statistically significant comparing infection induced by these two *Brucella* species.

**Figure 2.**
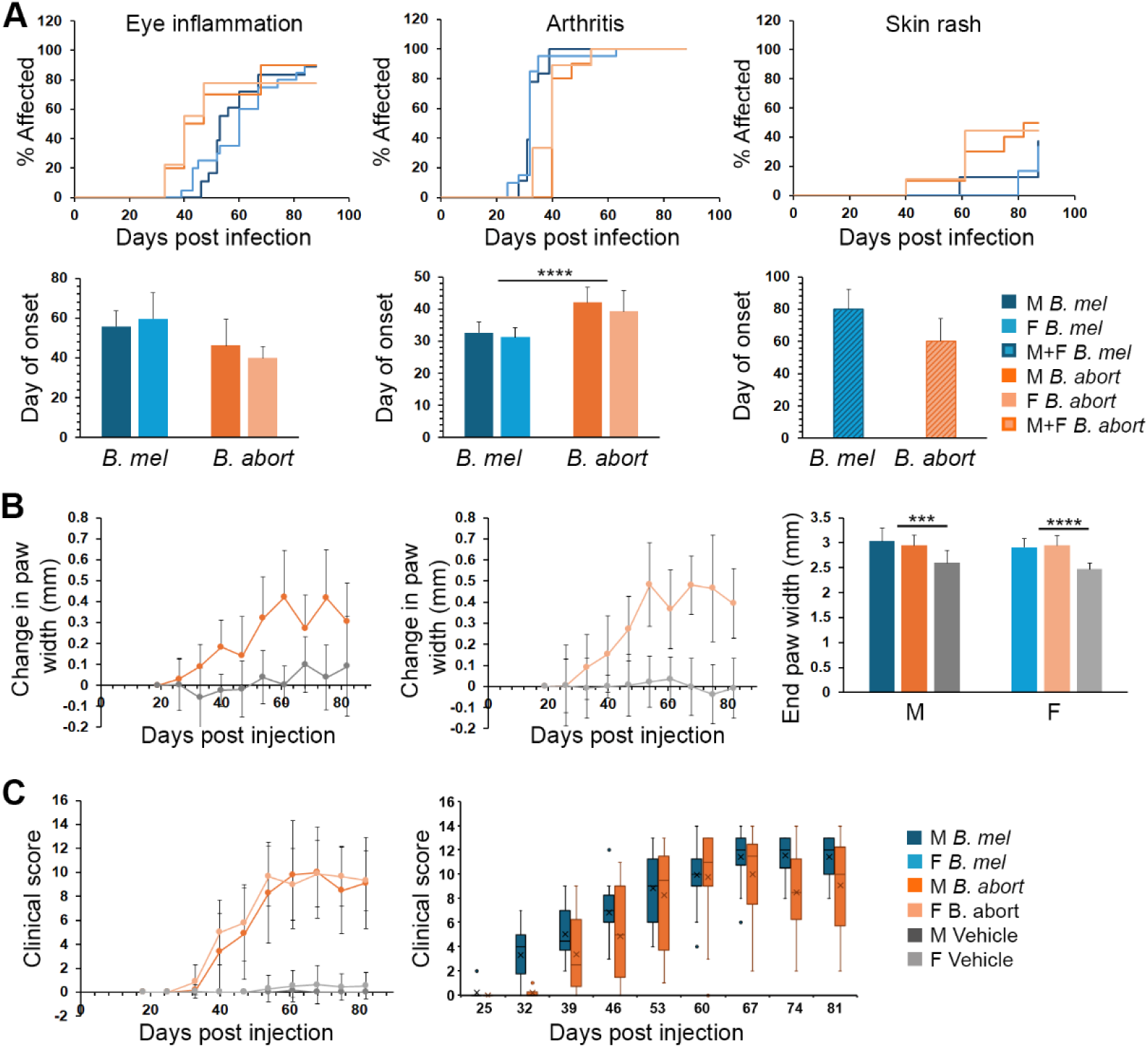
Comparison of *B. melitensis* and *B. abortus*-induced disease in SKG mice. Mice were injected with PBS (vehicle control, N=12 males (M) and 13 females (F)), 2×10^6^ *B. melitensis* 16M (*B. mel*, N=18 males and 19 females) or 2×10^6^ *B. abortus* 2308 (*B. abort*, N=10 males and 9 females) intraperitoneally (i.p.). A) Percent of mice (top row) that developed peri-ocular eye inflammation (left), arthritis (middle), and skin rash (right). Lower row is median day of onset for each symptom. Skin rash values are for combined males and females related to low prevalence. Day of rash onset represents average from 5 *B. melitensis* and 9 *B. abortus* mice. Error bars are S.D. B) Paw widths were measured by calipers, averaged as above and are vs. day 14-18 post-injection. Only infected animal paw widths significantly changed over time (p<0.001). End paw widths are at Day 81. C) Clinical scores over time in males and females infected with *B. abortus* (left) vs PBS vehicle control (p<0.03 after day 25 for infected vs. control animals). Right graph shows box whisker plots with average, median, 25^th^ and 75^th^ percentile, 10^th^ and 90^th^ (whiskers) and outlier points for *B. melitensis* vs. *B. abortus* infected males. ***p<0.005, ****p<0.001.

*B. neotomae*, a species found in wood rats and very rarely in humans, elicited even milder disease and revealed sex differences: No peri-ocular inflammation was observed in any *B. neotomae* infected mice. Arthritis penetrance was 40% in males vs 90% in females (p=0.15). Arthritis onset was significantly delayed in *B. neotomae* infected mice, as opposed to the more virulent *Brucella* strains (median onset of ∼7.5 weeks **Figure 3A**, p<0.001 vs. *B. melitensis*,). A couple of females developed paw skin rash, but no males. Changes in clinical score over time were significantly greater for female mice vs males (p<0.001), although individual time point comparisons were not statistically significant due to variability. In comparisons with *B. melitensis* infected mice, both *B. neotomae* males and females exhibited milder disease (p<0.001). In contrast to the females, clinical scores in *B. neotomae* infected males did not differ significantly vs controls (**Figure 3B**). Similarly, for change in paw widths over time, *B neotomae* infected males were mildly different than controls at individual times, whereas the female mice displayed greater differences, particularly later in the course (**Figure 3C**). End paw widths for males were no different than controls. To determine if a higher initial bacterial load would exacerbate disease, we infected mice with a log more *B. neotomae* (**Figure 3D and 3E**). The trends for penetrance were not significant. Females, but not males developed more severe arthritis by week 5 with the higher infectious dose (p<0.05). Interestingly, these mice still did not develop eye or skin disease, distinct from the *B. melitensis* mice.

**Figure 3.**
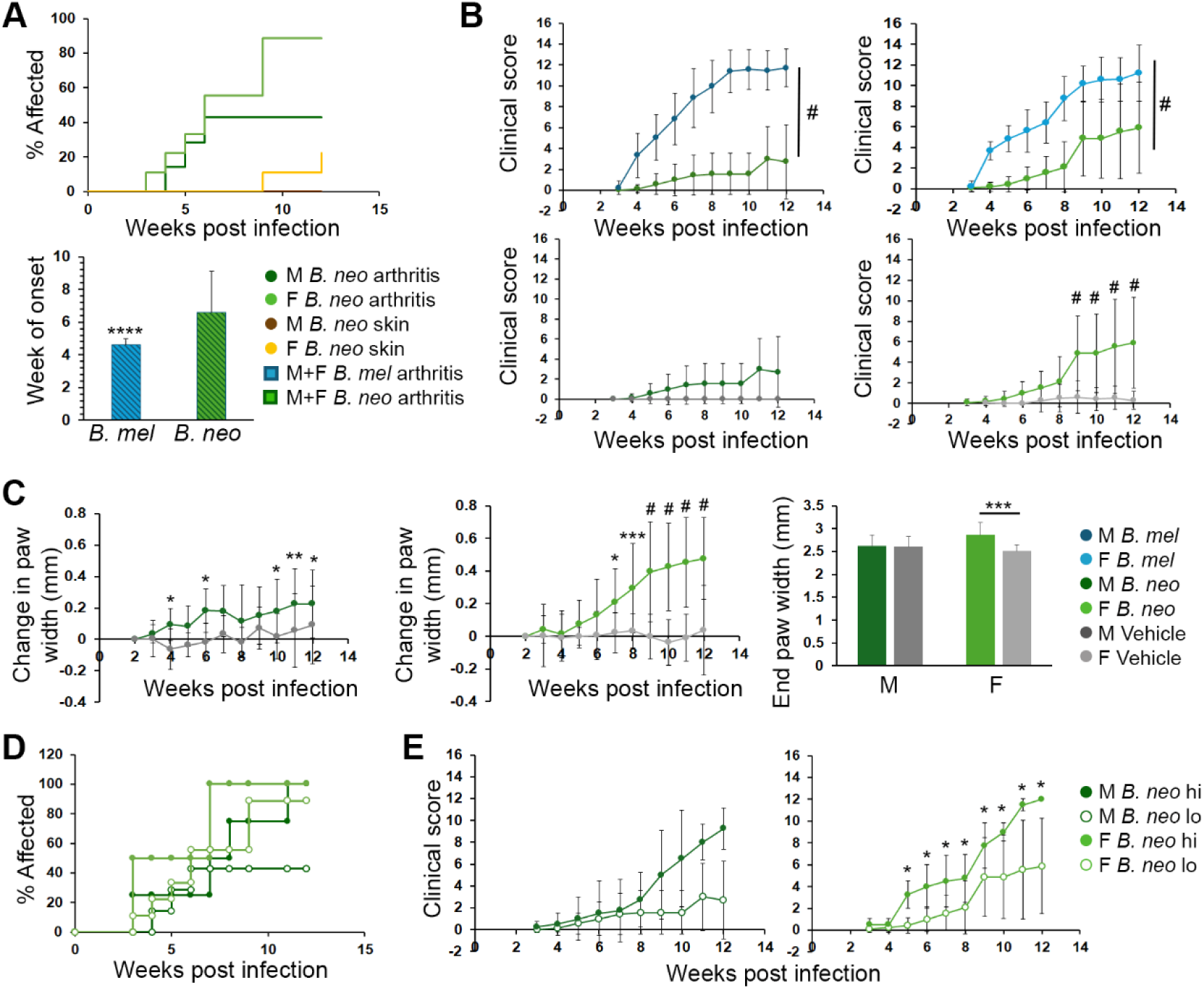
Disease onset, clinical scores and arthritis following *B neotomae* (*B. neo*) infection. SKG mice were infected with 2×10^6^ *B. neotomae* or *B. neotomae* containing the lux operon (N=7 males (M) and 9 females (F)). A) Percent of mice affected by arthritis and skin rash (top) and comparison with *B melitensis* for day of arthritis onset. B) Clinical scores over time with comparison to *B. melitensis* (top row) and PBS vehicle control (bottom row). In both sexes, *B. neotomae* mice exhibited milder disease by week 4 as compared to *B. melitensis* infected animals p<0.001. Males are on the left and females on the right. *B. neotomae* infected male clinical scores did not differ vs. controls. For females p<0.001 vs. controls over time and at individual time points by week 9. C) Changes in paw widths in mm vs week 2 for *B. neotomae* infected mice or vehicle control and end paw widths after 12 weeks infection. Both sexes showed significantly increased paw width changes over time vs controls (p<0.001 for rate of change). Average rate of change differed between males and females (p<0.001), although individual time point values were not significant between sexes. Individual time point pairwise comparisons are noted on the graphs. D) Percent of mice developing disease manifestation and E) Clinical scores for mice infected with either low (2×10^6^, open circles) or high dose (2×10^7^, closed circles) *B neotomae*. N=4 of each sex in the high dose groups with comparison to mice in 3A-3C. *P<0.05, **P<0.01, ***p<0.005, **** and #, p<0.001.

### Persistent *Brucella* infections in SKG spleen and paws

It was not clear if *Brucella*-induced arthritis in the SKG model is a purely “reactive process” to an infectious trigger or requires persistent infection. The dose response above suggested bacterial load was important. To further address this question, we used heat killed *Brucella*. Infection with heat killed *B. melitensis* did not induce obvious arthritis in comparison to live *Brucella*, although paw size subtly increased, suggesting the need for a viable infection rather than simple pathogen associated molecular pattern (PAMP) stimulation for significant arthritis (**Figure 4A**). However, some mild peri-ocular inflammation was noted starting at day 46 (**Figure 4B**) and there were no significant differences in rate of eye disease accrual or score. Heat-killed *B. melitensis* treated mice also failed to gain weight, exhibiting sawtooth fluctuations (**Figure 4C**). To examine live, but persistence-defective *Brucella*, we used a *B. melitensis* mutant lacking the Type IV secretion system (T4SS) encoded by the VirB operon (Δ*virB*). Preliminary data with these mutants suggests a similar response to the heat killed *Brucella* infection, with no significant arthritis and mild eye disease (**Figure S2**). *Brucella* have been reported to persist in BALB/c spleens beyond 90 days, raising the question of whether the SKG mice were also impaired in clearance. In SKG mouse spleens harvested at 12 weeks, we observed differences between species, with *B. melitensis* showing the highest splenic CFU, *B. abortus* intermediate with about a log less, and *B neotomae* with very low levels of splenic CFU (**Figure 4D**). Spleens from mice administered heat killed *Brucella* and Δ*virB* were negative for CFU at the end of the experiment, as expected. Interestingly, splenic CFU from live *Brucella* infected animals correlated moderately well with 12-week clinical score (ρ=0.79, **Figure 4E**), with this correlation mainly driven by differences in *Brucella* species, suggesting a relationship between the severity of systemic infection and arthritis.

**Figure 4.**
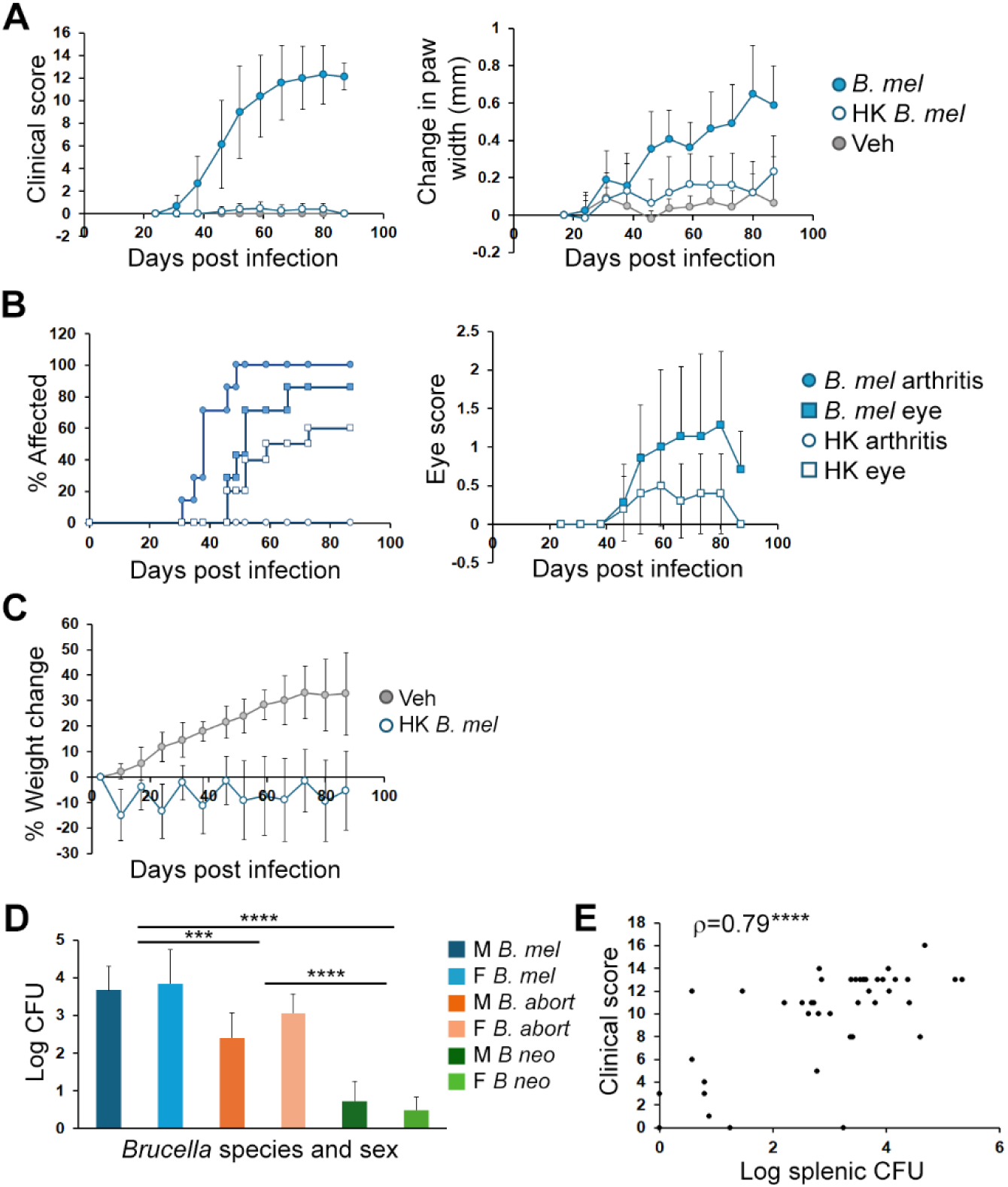
Contributions of persistent colonization to disease severity. A-C) Mice were injected i.p. with PBS vehicle control (Veh, 4 mice), live *B. melitensis* (*B. mel*, 7 mice), or heat killed *B. melitensis* (HK, 10 mice). Males and females were combined. A) Joint and eye disease was scored over time (left) and changes in paw widths in mm vs 2 weeks (right). Pairwise comparisons of *B. melitensis* and heat killed *Brucella* clinical scores were p<0.03 by 4 weeks and change in score over time was p<0.001. Paw widths for heat killed *Brucella* changed significantly compared to both control animals and to viable *B. melitensis* (p<0.019 for all pairwise comparisons). B) Percent of mice developing disease manifestations (left) and eye disease score (0-2, right). Accrual over time only differed for arthritis (p<0.001). Average eye scores did not significantly differ. C) Percent change in weight over time as compared to week 0. P<0.001 for weight gain in controls only. D) At 12 weeks following infection with *B. melitensis* (*B. mel*, M= 16 males, F= 11 females), B. *abortus* (*B. abort*, 4 each sex), *B. neotomae* (*B. neo*, 4 each sex), or heat killed *Brucella* (10) splenic CFU were enumerated. E) Correlation between final clinical score at 12 weeks and splenic CFU for the mice in (D). For the N=10 heat killed *Brucella*, all CFU and final clinical scores were 0 (set to 0.1 for purposes of graphing). Spearman’s rank correlation coefficient is 0.79, p<0.001. ***p<0.005, ****p<0.001.

Previous studies using in vivo imaging (IVIS) and our histology sections (**Figure 1F**) suggested *Brucella* colonize the paws during infection. To determine if SKG paws contained viable *Brucella*, articular tissue was dissected and evaluated for CFU. Paw tissue from *B. melitensis* infected mice reliably grew detectable CFU at 12 weeks (**Figure 5A**). Since the amount of tissue recovered from swollen paws was variable, we examined normalized gene expression for several UPR genes (*Hspa5*/BiP, *Ddit3*/CHOP and spliced *Xbp1* (marker of active UPR)) and UPR-regulated cytokines (*Tnf, Ifnb1* and *Il23a*) to gauge relative degree of infection. Indeed, normalized gene expression from paw tissue revealed greater levels of cytokines, and a trend for *Hspa5* and *Ddit3*, suggesting the paws had a relatively greater burden of infection compared to spleen (**Figure 5B**).

**Figure 5.**
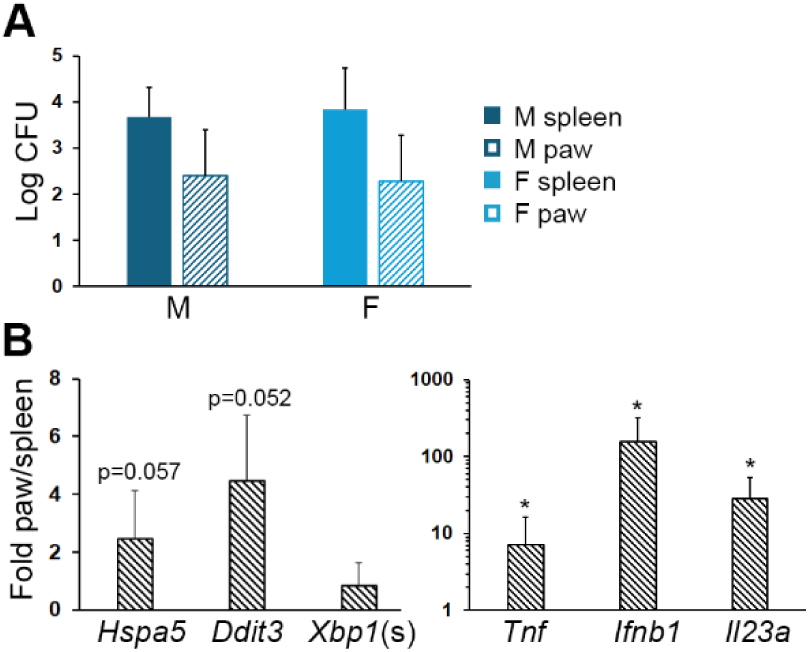
Paw CFU and gene expression. Male (M) or Female (F) mice were infected with 2×10^6^ *B. melitensis* for 12 weeks prior to harvest of spleens and paw tissue. A) Log CFU from spleens and paws. Results are from 15 male spleen, 5 male mouse paws, 11 female spleen and 4 female mouse paws. B) Spleens and paws were harvested for RNA and gene expression determined by quantitative PCR with normalization to the housekeeping gene 18S rRNA. Fold mRNA is the average ratio of paw to spleen gene expression within each mouse from 3 males and 4 females. *p<0.05 for the ratio being different from 1.

To further clarify the anatomic distribution of *Brucella* in infected SKG mice, we *used in vivo* imaging (IVIS) to track luminescent *B. neotomae*. Expression of the Lux operon requires ATP production and thus viable bacteria, and the excellent correlation with CFU has been previously described^9^. *Brucella* were present in the paws by one-week post-infection and remained there for the duration of the experiment, even when the spleen and liver did not exhibit any luminescent signal (**Figure 6A**). Week 1 appeared to be the height of paw colonization by quantification. Other consistent luminescent hot spots included the neck and genital areas. Paw luminescence was quantified as in **Figure 6B** and after the first 2 weeks, exhibited relatively constant levels over the 12-week experiment, with most paw luminescence gently sloping off during the course of infection. Average luminescence in the paws was not significantly different for infection with 10^6^ vs 10^7^ *Brucella*, despite some differences in clinical severity (**Figure 3E**), and thus results were combined. Over time, infected females had greater paw luminescence vs males, though around week 9, the two curves started to converge (**Figure 6C**). Overall, there was a positive correlation for luminescence and clinical score areas under the curve, suggesting an impact of infectious paw burden over time (**Figure 6D**). However, tracking both luminescence and clinical scores at individual time points and in individual mice revealed a different story: there was no arthritis apparent early during peak luminescence and clinical scores increased even as luminescence levels tapered (**Figure 6E**). Thus, arthritis severity at any given time did not directly reflect the amount of live *Brucella* in the paws.

**Figure 6.**
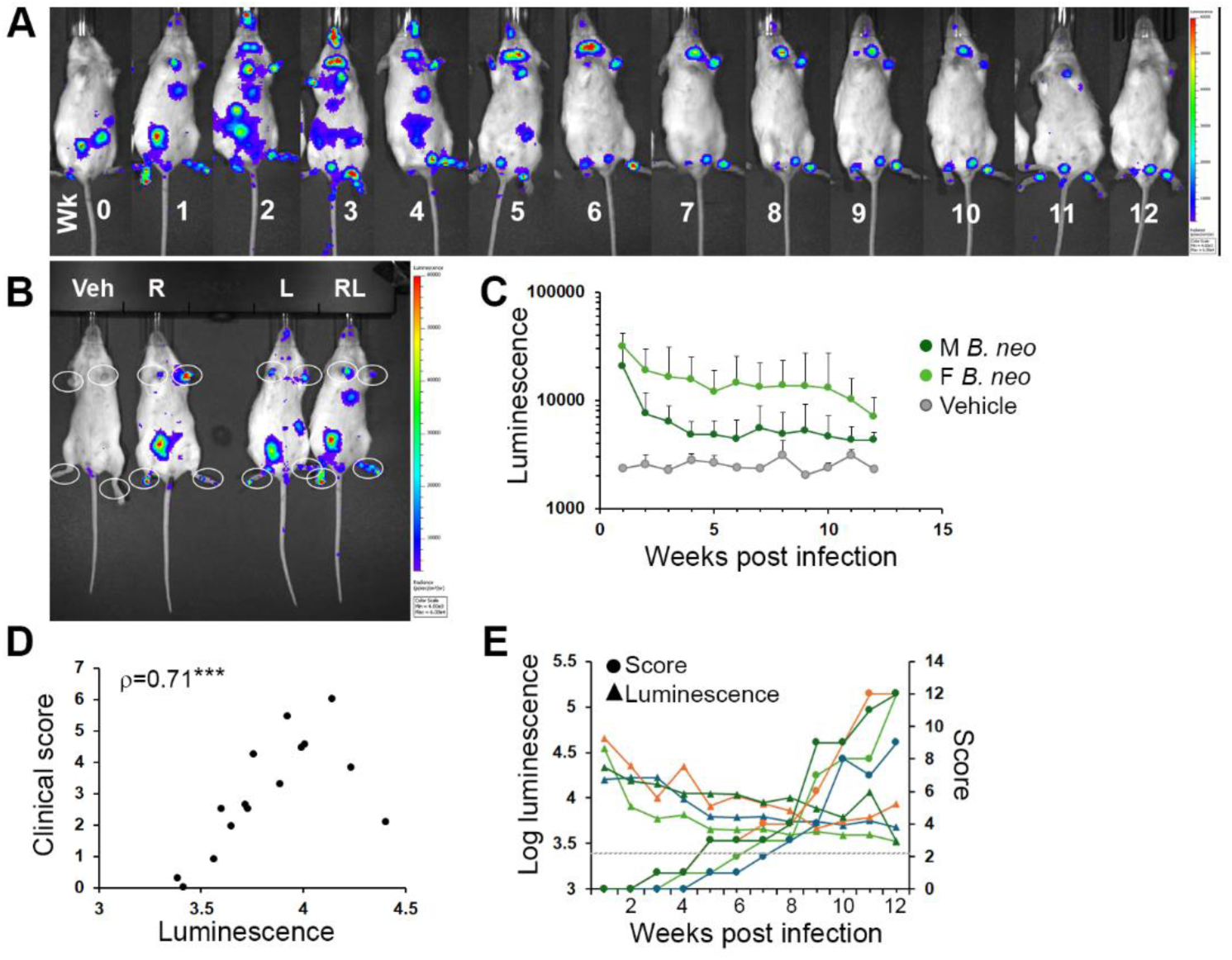
Distribution of active infection and paw colonization in B. neotomae infected mice over time. Mice were infected intraperitoneally with 2×10^6^ or 2×10^7^ *B. neotomae* expressing the Lux operon. A) Sample mouse luminescence over time (wk = weeks) showing anatomic distribution. B) For quantification, regions of interest were drawn around all 4 paws and luminescence averaged per mouse. Example image is from mice 1-week post-infection with *B. neotomae* (R, L and RL) or post injection of PBS vehicle control (Veh, left-most mouse). C) Luminescence data from 7 females, 6 males and 2 controls (1 of each sex). For analysis, average luminescence was log10 transformed. Luminescence and clinical score area under the curve was divided by the time range to give an “average value” over time. Control mice vs. infected p=0.019. Infected males vs females p=0.004. D) Correlation between average transformed luminescence and average clinical score (area under the curve), ρ=0.71, p=0.003. E) Log luminescence (triangles) is plotted vs corresponding clinical scores (circles of same color) for 4 sample females showing values at individual time points. Gray dashed line is average background luminescence.

### Use of the SKG model to probe attenuated *Brucella* mutants

The disease score differences seen with the various *Brucella* species suggested this mouse model might be useful for evaluating the consequences of *Brucella* mutations for clinically relevant outcomes. The T4SS is required for *Brucella* secretion of effectors that modulate the macrophage intracellular milieu. One of these effectors, TcpB (also known as BtpA) has been reported to alter the inflammatory milieu *in vivo* without significant effect on colonization, providing another opportunity to assess the reactive component of disease. TcpB is also one of several *Brucella* effectors that support a pro-inflammatory endoplasmic reticulum response known as the Unfolded Protein Response (UPR). Mice infected with a TcpB deletion mutant (Δ*tcpB*) *B. melitensis* consistently displayed lower clinical scores throughout the 12-week period as compared to the wild type *B. melitensis*, as well as decreased paw width changes (**Figure 7A**). Also, arthritis onset was a week later for the TcpB mutant infected mice and no TcpB mutant-infected mice developed peri-ocular inflammation or skin rash (**Figure 7B**). Splenic CFU was more variable in the ΔTcpB mutant-infected mice, but not statistically different from the parental *B. melitensis* wild type (**Figure 7C**). There was a non-significant trend towards decreased *Ddit3* (CHOP) expression in spleens and paws and average UPR gene expression in the paws of the Δ*tcpB* mutant-infected mice (**Figures 7D**).

**Figure 7.**
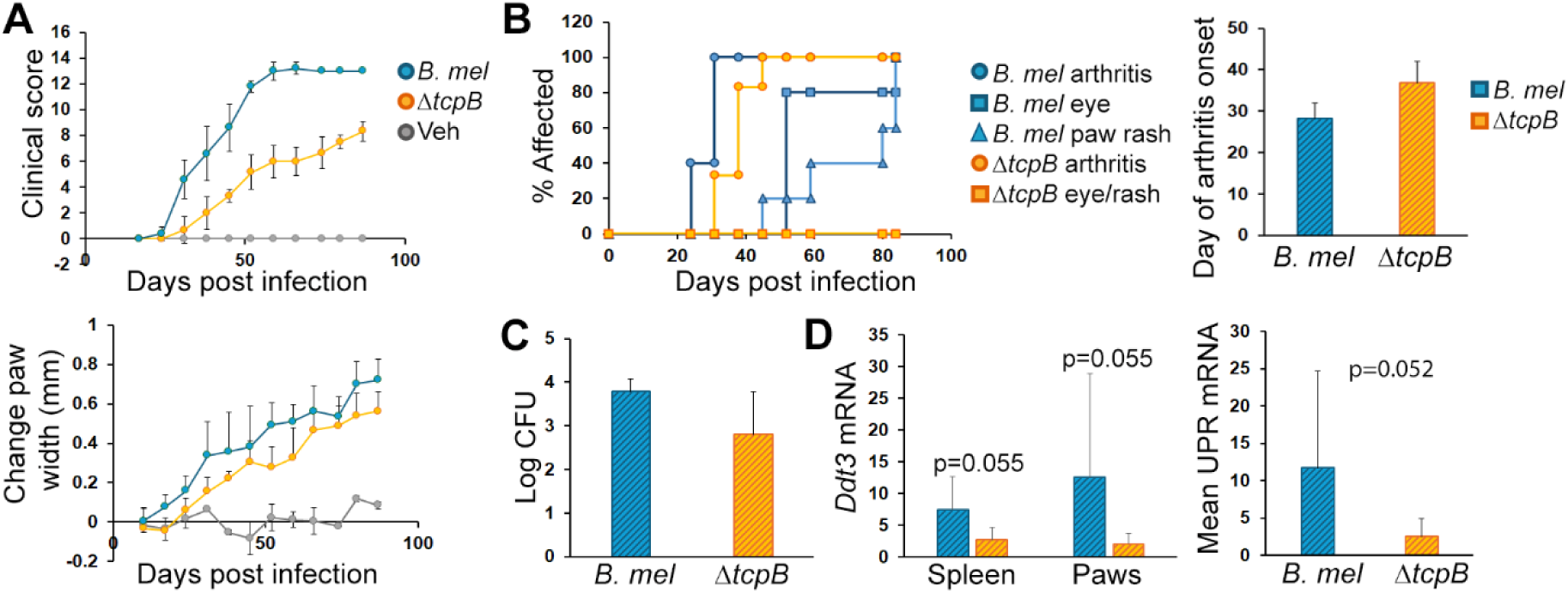
Milder disease with Δ*tcpB Brucella* mutant. Mice were infected i.p. with 2×10^6^ *B. melitensis* (*B. mel*, 3 males (M) and 2 females (F)), 2×10^6^ Δ*tcpB* mutant B. melitensis (3 M and 3 F), or PBS vehicle control (Veh, 1 M and 1 FA). A) Mice were scored weekly for 12 weeks. *B. mel* vs Δ*tcpB* over time and in pairwise comparisons after day 25 p<0.001. Paw widths were measured by calipers weekly and averaged per mouse. All paw widths in infected mice change over time (p<0.001), and comparisons of linear trends between wild type, mutant *Brucella* and controls are all p<0.002. B) Percent of mice developing arthritis, peri-ocular eye inflammation and psoriatic paw rash. Arthritis onset was later in Δ*tcpB* vs wild type, p=0.02. C) At 12 weeks, spleens were harvested, and *Brucella* quantitated by colony forming unit (CFU). CFU are not significantly different. D) Spleens and paws were harvested for RNA, and mRNA quantitated by qPCR with normalization to 18S rRNA. Left is *Ddit3* expression and right is the combined averages of *Xpb1*(s), *Ddit3* and *Hspa5* gene expression in paw synovial tissue.

## Discussion

Herein, we describe a fully penetrant, easily scored clinical model of *Brucella* induced spondyloarthritis in SKG mice with cardinal features of large joint arthritis, peri-ocular inflammation and scaly paw rash. Except for some mild breakthrough disease in two females when the colony was first getting established, non-infected controls have presented a clean comparator group. The model has enough dynamic range to reveal differences between wild type and mutant *Brucella*, as well as between different *Brucella* species. The more severe disease in *B. neotomae* infected-females vs males is intriguing. It is not yet clear whether this difference was revealed because of milder infection by *B. neotomae* (as evidenced by the lower splenic CFU vs *B. melitensis*) or whether this is a *Brucella* species-specific finding. The lack of eye and skin disease with higher dose *B. neotomae* support the notion of species-specific inflammation. Results from the Δ*tcpB* and Δ*virB* mutants suggest this model will be useful for investigating the genetic and molecular correlates of bacterial virulence *in vivo*.

The presence of splenic and paw CFU at 12 weeks, particularly in the *B. melitensis* and *B. abortus* infected mice and paw luminescence in the *B. neotomae* females revealed active persistent infection. Moreover, during *Brucella* infection, there was a moderately positive correlation between clinical score and splenic CFU, particularly with the higher CFU infections. *B. neotomae* infected females developed moderate to severe arthritis despite low splenic CFU, thus paw colonization may be particularly important. The absence of significant arthritic disease with heat killed bacteria support the idea that persistent infection is required for arthritis and suggests PAMPS alone are not sufficient to elicit all disease manifestations. However, the mild eye inflammation and failure to gain weight suggest the PAMP load is still having an adverse effect on the mice. On the other hand, the lack of strict correlation between luminescence and arthritis in the paws at individual time points suggests the arthritis does not simply reflect degree of local infection at any given time but requires the development of an additional host inflammatory response, or a “reactive” component. The results from the *Brucella* mutant infections also support the idea of a reactive component, as we confirmed the ΔTcpB mutant consistently displayed an intermediate phenotype for most of the experiment duration without significant effects on splenic CFU.

It is not entirely clear whether SKG infection represents an apt model of human brucellosis or should be considered more of a non-specific infection-triggered spondyloarthritis model. The most common osteoarticular complications of brucellosis in humans consist of large joint arthritis (particularly knees), sacroiliitis and spondylitis. Disease induced by the different *Brucella* species correlated somewhat with reported virulence in humans in that *B. melitensis* generally leads to the most prevalent and severe human infections and *B. neotomae* rarely causes human disease – although *B. neotomae* has been rarely reported in human neurobrucellosis. Axial disease was not clearly evident in *B. neotomae* infected mice by IVIS, but we do not have the facilities to examine this question using luminescent *B melitensis* (a BSL3 select agent). The SKG mice did have a 3–4-week delay in developing overt signs of arthritis, so onset of clinical disease is during the “chronic” phase of *Brucella* infection. In humans, osteoarticular disease takes several weeks to develop, with >60% occurring after 3 months. Also, up to 21% of humans with Brucellosis develop ocular complications, most often uveitis. Interestingly, prevalence of uveitis in ankylosing spondylitis is ∼25%. Skin disease in humans with Brucellosis is more uncommon, present in 1-14%. However, cutaneous involvement in humans is more often characterized by maculopapular skin eruption, rather than scaly psoriasis-like rash.

The phenotype of *Brucella* infected mice differs from that of curdlan injected mice in several ways: Onset of arthritis is rapid in curdlan injected females, occurring 7 days post-injection. Male mice have delayed arthritis onset and less severe scores, although 100% eventually get arthritis by day 35. Curdlan injected mice develop “dactylitis” or fusiform digital swelling, which was absent in our model. Only about 25% get eye disease, which consists of posterior uveitis and peri-ocular inflammation as seen in our model, though with greater penetrance in our studies. Interestingly, new bone formation about the wrist is also seen following curdlan, along with severe erosions. A prominent feature of the curdlan-induced inflammatory disease is small bowel inflammation (ileitis) occurring in 50-60% by 10-12 weeks. In our model, there was no inflammation evident in the gut grossly or by histologic section. However, it is possible that we may not have looked far enough out, as experiments were ended at 12 weeks. Curdlan-driven disease is both gut microbiome and Th17-dependent.

*Chlamydia* infection of SKG mice produces a different phenotype as well. Male-female differences are more pronounced in this model than what we observed. Following genitourinary infection, 85% of females get arthritis but male mice are spared. However, following intranasal infection, 50% of male mice get milder arthritis. Arthritis severity is asymmetric, unlike the experience with *Brucella*, where all paws were affected and to a similar extent. *Chlamydia* infected mice also do not get dactylitis, as in our data. Compared to females, male mice infected with *Chlamydia* are much more prone to developing eye inflammation. Neither sex demonstrates ileitis. Similarly to our results with *Brucella*, disease severity correlated with infectious dose and degree of genitourinary inflammation. The authors suggested reactive inflammatory disease resulted from insufficient sterilization of infection by SKG mice, as there was decreased IFN-γ and IL-17 production, but excessive TNF-α.

Much work remains to establish the pathogenetic mechanisms underlying *Brucella* induced spondyloarthritis disease in this robust model. Moreover, the mechanisms may differ by site of involvement (e.g. eyes vs joint). On the host side, it will be interesting to determine how the altered T cell repertoire in the SKG modulates control and anatomic distribution of *Brucella* as compared to the parental BALB/c strain. The host UPR has been implicated in rheumatologic spondyloarthritis. However, a recent report, wherein CHOP-deficient HLA-B27 transgenic rats actually developed similar or worse gut disease despite the lack of CHOP-dependent IL-23, suggests the relationship between UPR and inflammation in spondyloarthritis will require further elucidation. *Brucella* induces a full UPR in infected macrophages, and thus this model presents an incredible opportunity for further dissection. Further investigation of the male-female differences with *B neotomae* infection may offer insight into the effect of sex on anti-bacterial immune responses. The capacity to observe gradations in mouse disease severity as well as alterations in onset and sites of inflammatory involvement offers the potential for defining bacterial correlates contributing to virulence *in vivo*. In conclusion the SKG mouse model should provide remarkable new opportunities for understanding the connections between *Brucella* infection and spondyloarthritis features.

## MATERIALS AND METHODS

### Zap70^SKG/SKG^ (SKG) mice and infection protocol

The SKG mouse line, first described by Sakaguchi and colleagues, carries a homozygous G-to-T substitution at nucleotide 489 in the *Zap70* gene on the BALB/c genetic background ^20^. SKG mice were kindly donated to our laboratory by Drs. David Riches and Elizabeth F. Redente of National Jewish Health, Denver CO. The mice were rederived using pathogen-free recipients in the Genome Editing Facility of the University of Wisconsin-Madison’s (UW) Biomedical Research Model Services (BRMS). Animal research described in this study followed protocols approved by the UW Institutional Animal Care and Use Committee. The UW-Madison is an AAALAC approved institution. For infection, *Brucella* were grown to late log to stationary phase, quantified, and diluted to 1 x 10^6^/100 µl in PBS. Each mouse was given 200 µl intra-peritoneally. Control animals received injections of vehicle (PBS) only. *Brucella* were heat-killed (HK) through incubation at 56°C for 1 hour.

### Brucella strains

All bacteria used in this study were cultured in Brain Heart Infusion Broth/Agar (DOT Scientific). Antibiotics, when required, were added at the following concentrations: carbenicillin, 100 mg/L; kanamycin 100 mg/L. Quantification of bacteria was from established growth curves through Optical Density measurements at 600 nm using a UV-Visible Spectrophotometer (Thermo Scientific). *Brucella* abortus S2308 and *Brucella melitensis* 16M were from University of Wisconsin archived stocks. *Brucella neotomae* was purchased from the American Type Culture Collection (ATCC). The virB operon knock-out mutant of *B. melitensis* (Δ*virB*) was generated as described ^21^ using the plasmid pAVT1.4 (generously provided by Dr. Renee Tsolis). The *tcpB* (*btpA*) knock-out mutant of *B. melitensis* (Δ*tcpB*) was generated as described ^22^. Bioluminescent *B. neotomae* expressing the luxCDABE operon was generated as previously described ^9^. All experiments with select agent *Brucella* strains were performed in a Biosafety Level 3 (or Biosafety Level 2 for *B. neotomae* only) facility in compliance with the Federal Select Agent Program (FSAP) in accordance with standard operating procedures approved by the University of Wisconsin-Madison Institutional Biosafety Committee.

### Clinical disease assessment

Arthritis assessment began four days after injections and continued weekly for 12 weeks. A thickness gage (Mitutoyo) was used to measure paw thickness at wrist joint of the front paws and ankle joints of hind paws. Measurements were recorded in millimeters. Paws were individually scored concordant with swelling and erythema, along with overall periocular inflammation as follows: paw swelling of 0 (none), 1 (mild), 2 (moderate), 3 (severe); eye inflammation of 0 (none), 1 (mild), 2 (marked). Peri-ocular inflammation was typically bilateral. The clinical score was a sum of arthritis score in all 4 paws plus periocular disease score (maximum of 14). For percent affected, the first day a symptom was observed was used for generating the Kaplan Meier survival curves. Unscored observations that were recorded (yes/no) included diarrhea, tail bone swelling, rough skin, and scaly psoriatic paw rash. Mouse weight was recorded weekly.

### Spleen and paw synovia tissue isolation

Mice were euthanized by carbon dioxide and rinsed thoroughly with 100% Ethanol. Spleens were placed in a 12 ml screw cap tube containing 3 ml MACS buffer (PBS, 2mM EDTA, 0.5% BSA) on ice. Mouse limbs were then dissected with scissors. Skin on each limb was peeled back and removed. Microsurgery scissors (FST 1502310) were used to cut remaining skin away from foot and paw. These scissors were then used to cut synovia from ankles, wrists, and paws and synovia were placed in a MACS buffer-containing tube as with the spleens above. Synovia from all 4 limbs were combined in the same tube. Tissues were then dissociated using the gentleMACS™ system (Miltenybiotec) using C tubes and following the manufacturer’s suggested protocol. After dissociation, tissue was divided equally (1.5 ml each) into two 1.7 ml microcentrifuge tubes. Samples were then pelleted in a microcentrifuge for 5 min at 400 x g.

For Colony Forming Unit (CFU) assay, 1 ml lysis buffer (dH_2_O + 0.1% Triton X-100 (Sigma)) was added to one of the tubes and thoroughly vortexed (at least 20 seconds). For quantitative PCR (qPCR) assay, 0.5 ml RNAzol RT reagent (Molecular Research Center, Inc.) was added to the second tube and likewise thoroughly vortexed (at least 20 seconds). Samples for RNA isolation were then frozen at −80°C for later processing while samples for CFU assay were processed immediately.

### Colony Forming Units (CFU) and quantitative PCR (qPCR)

A modified most probable number (MPN) approach was used to enumerate *Brucella* in mouse spleen and synovia samples using 10-fold serial dilutions ^23^. Eight replicates were plated onto a square petri dish (100 mm x 100 mm) with grid (FisherScientific) containing BHI agar. CFU were enumerated after 4 days growth.

Samples for total RNA isolation were thawed in RNAzol RT (MRC) and processed following the manufacturer’s protocol. RNA quantity and quality was determined using a NanoDrop One spectrophotometer (ThermoFisher). RNA was reverse transcribed using random primers (Superscript III; Invitrogen) following the manufacturer’s recommended protocol and cDNA quantified on a StepOnePlus Real-Time PCR System (ABI) normalized to 18S rRNA using the standard ΔCt/ΔCt method. The mouse primers used in this study were designed using IDT’s online primer design tool and mixed with template in PowerUp™SYBR™Green master mix (ThermoFisher) following the manufacturer’s recommended protocol. Primer sequences: 18S rRNA: F: GGACACGGACAGGATTGACAG; R: ATCGCTCCACCAACTAAGAACG *Hspa5*: F: AGGATGCGGACATTGAAGAC; R: AGGTGAAGATTCCAATTACATTCG *Ddit3*: F: CATCACCTCCTGTCTGTCTC; R: AGCCCTCTCCTGGTCTAC *sXbp1*: F: GAGTCCGCAGCAGGTG; R: GTGTCAGAGTCCATGGGA *Ifnb1*: F: GGCATCAACTGACAGGTCTT; R: ACTCATGAAGTACAACAGCTACG *Il23a*: F: ACAAGGACTCAAGGACAACAG; R: TGAAGATGTCAGAGTCAAGCAG *Tnf*: F: TCTTTGAGATCCATGCCGTTG; R: AGACCCTCACACTCAGATCA

### In Vivo Imaging, Quantification, and Micro-Computed Tomography (CT)

*Brucella neotomae* expressing the lux operon (luxCDABE) as an autonomous bioluminescent reporter was used to visualize bacterial dissemination and tissue localization within live SKG mice using an In Vivo Imaging System (IVIS Spectrum, Perkin Elmer). *B. neotomae* expressing lux (B. neo/lux) was generated using transposon transformation as previously described ^9^. SKG mice infected with *B. neo*-lux were housed and imaged at the University of Wisconsin-Madison Small Animal Imaging and Radiotherapy Facility (SAIRF) Luminescence was quantitated using Living Image® Software (PerkinElmer). Since bioluminescence correlates with bacterial numbers ^22,24^, Regions of Interest (ROI) over the paws were used to quantitate dissemination and persistence of *Brucella*. In vivo imaging and arthritis assessment (as described above) of the *B. neo*-lux mice was performed weekly for 12 weeks.

Ex vivo µCT was performed at the University of Wisconsin-Madison Small Animal Imaging & Radiotherapy Facility (SAIRF) using the MILabs U-SPECT/CT^UHR^ (ultra high-resolution) system. Samples were prepared as follows: Mice were euthanized, and limbs were dissected. Skin was removed from each limb. Limbs were then placed in 12 ml conical screw-cap tubes (CELLTREAT) partially filled with EMA fixative (EtOH, methanol, acetic acid; 3:1:1). Then, gauze was inserted to keep limb in place, and tube filled to top and capped. After 24hrs, samples were scanned and images prepared by SAIRF staff (Dr. Justin Jeffries).

### Histological Preparation and Analyses

Sample preparation of front and hind paws for histological staining followed the standardized microscopic arthritis scoring of histological sections (SMASH) recommendations ^25^. Briefly, paws were cut 2-3 mm above the ankles or wrists and the skin was removed. Samples were placed in 12 ml screw cap centrifuge tubes and filled with Decalcifier I (Surgipath) for fixation and decalcification. After 24h, samples were sent to the Comparative Pathology Lab (CPL) of the UW-Madison where histology staff prepared H&E-stained slides. Slides were imaged and evaluated for arthritis by a Pathologist (Dr. Thomas Warner) of the Department of Pathology and Laboratory Medicine.

### Statistical Analysis

Time to onset of disease manifestation was estimated using Kaplan-Meier methods and compared among treatment groups using log-rank tests. Poisson regression (with offset equal to the logarithm of follow-up time) was used to estimate occurrence rates for each treatment group and manifestation (eye, arthritis, skin), with cluster-robust standard errors used to account for multiple manifestations within the same mouse. Linear mixed-effect models were used to evaluate weight and paw width (change) over time. These models were also used to determine differences between treatment groups at individual time points, with p-values adjusted for multiple comparisons to control the false discovery rate (FDR). Clinical scores were compared between groups of interest at individual time points using rank-based methods (Wilcoxon rank-sum test) and p-values again adjusted (FDR method) for multiple testing. Scores were also summarized by computing the area under each animal’s score profile over time; ordinary linear models were used to compare the resulting distributions of areas among groups of interest (defined by sex and/or treatment) or to determine whether there was a correlation (Spearman’s rho) with similarly computed areas over time for luminescence. Ordinary linear models were used to analyze (log-transformed) CFU abundance and UPR or cytokine gene expression for associations with treatment or sex. All p-values were 2-sided unless expressly stated. Analyses were performed using R (v. 4.4.0; https://cran.r-project.org/). Error bars represent the means ± STD DEV. Statistical significance is indicated in the figures (* p<0.05, ** p<0.01, *** p<0.005, **** p<0.001) and figure legends.

## ACKNOWLEGEMENTS

The authors would like to acknowledge the Cancer Center Support Grant: NCI P30 CA014520, University of Wisconsin Small Animal Imaging & Radiotherapy Facility and NIH S10OD028670-01, and NIH 2R01AI116453-06 (PI: Oliveira) for supporting this work.

The authors are grateful to Drs. Elizabeth Redente and David Riches of National Jewish Health for providing the initial SKG breeding mice. The authors thank the University of Wisconsin School of Medicine and Public Health Biomedical Research Model Services shared resource for the use of its facilities and their Research Services team for expertise including breeding and specialized techniques. The authors also thank Dr. Renee Tsolis for the *vir*B operon knock-out plasmids and Dr. Suzana Salcedo for her discussion and support.

## Contact for Reagent and Resource Sharing

Further information and requests for resources and reagents should be directed to and will be fulfilled by the lead contact, Judith A. Smith

---

**Supplemental Figure S1:**
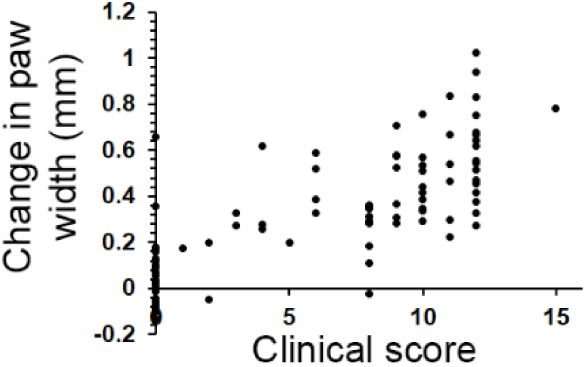
Correlation between paw width measurements and clinical scores. Summary of mice infected with *B. melitensis, B. abortus, B. neotomae*, or injected with PBS vehicle control (N=39, 19, 16 and 25 respectively). Data points are end-experiment change in paw width vs. day 14-18 (in mm) and final clinical scores at days 81-88. Spearman’s correlation coefficient ∼ 0.76, p<0.001. There is also a correlation with final paw width (ρ ∼ 0.69, p<0.001). The significance in correlation is driven largely by differences in infection treatment group.

**Supplemental Figure S2:**
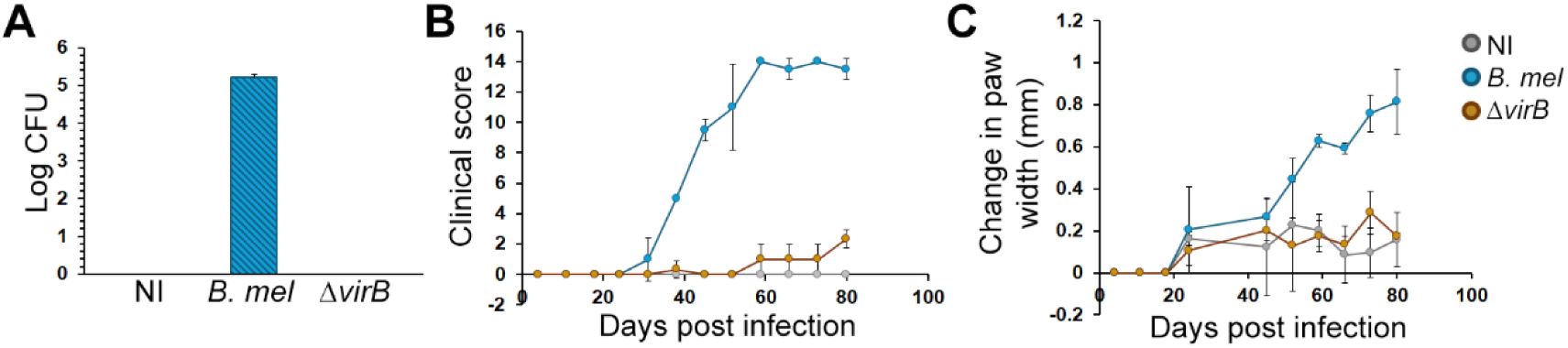
The Type IV secretion system is required for *Brucella* persistence and the development of arthritis. Mice were injected with PBS (NI, non-infected, gray), wild type *B. melitensis* (*B. mel*, blue) or a mutant lacking the VirB operon encoding the Type IV secretion system (Δ*virB*, brown). Male and female mice were combined, with 3 Δ*virB* and 2 each of the *B. melitensis* and PBS controls. A) Splenic CFU at 12 weeks. B) Clinical scores over time. Note, the Δ*virB* mice developed mild peri-ocular inflammation but not arthritis. *B. melitensis* score trajectory was significant vs Δ*virB* (p<0.001), and Δ*virB* was not significant vs control (p=0.093). C) Changes in paw widths over time. Data points are on days of measurement by calipers and changes in millimeters are vs. day 18 (set=0). All animal paw widths increase over time (p<0.01). *B. melitensis* paw widths exhibit greater change vs. Δ*virB* over time (p<0.001). Δ*virB* paw width changes do not differ significantly from PBS controls.

## References

1. Pappas G, Papadimitriou P, Akritidis N, Christou L, Tsianos EV. The new global map of human brucellosis. Lancet Infect Dis 2006;6:91–9.

2. Rossetti CA, Arenas-Gamboa AM, Maurizio E. Caprine brucellosis: A historically neglected disease with significant impact on public health. PLoS Negl Trop Dis 2017;11:e0005692. PMC5560528

3. Pappas G, Akritidis N, Bosilkovski M, Tsianos E. Brucellosis. N Engl J Med 2005;352:2325–36.

4. Franco MP, Mulder M, Gilman RH, Smits HL. Human brucellosis. Lancet Infect Dis 2007;7:775–86.

5. Esmaeilnejad-Ganji SM, Esmaeilnejad-Ganji SMR. Osteoarticular manifestations of human brucellosis: A review. World J Orthop 2019;10:54–62. PMC6379739

6. Giambartolomei GH, Arriola Benitez PC, Delpino MV. Brucella and Osteoarticular Cell Activation: Partners in Crime. Frontiers in microbiology 2017;8:256. PMC5316522

7. Corbel MJ. Brucellosis: an overview. Emerging infectious diseases 1997;3:213–21. 2627605

8. Olsen SC, Palmer MV. Advancement of knowledge of Brucella over the past 50 years. Veterinary pathology 2014;51:1076–89.

9. Rajashekara G, Glover DA, Krepps M, Splitter GA. Temporal analysis of pathogenic events in virulent and avirulent Brucella melitensis infections. Cell Microbiol 2005;7:1459–73.

10. Durward M, Radhakrishnan G, Harms J, Bareiss C, Magnani D, Splitter GA. Active evasion of CTL mediated killing and low quality responding CD8+ T cells contribute to persistence of brucellosis. PLoS One 2012;7:e34925. 3338818

11. Magnani DM, Lyons ET, Forde TS, Shekhani MT, Adarichev VA, Splitter GA. Osteoarticular tissue infection and development of skeletal pathology in murine brucellosis. Dis Model Mech 2013;6:811–8. PMC3634663

12. Skyberg JA, Thornburg T, Kochetkova I, Layton W, Callis G, Rollins MF, Riccardi C, Becker T, Golden S, Pascual DW. IFN-gamma-deficient mice develop IL-1-dependent cutaneous and musculoskeletal inflammation during experimental brucellosis. J Leukoc Biol 2012;92:375–87. 3395417

13. Khalaf OH, Chaki SP, Garcia-Gonzalez DG, Ficht TA, Arenas-Gamboa AM. The NOD-scid IL2rgamma(null) Mouse Model Is Suitable for the Study of Osteoarticular Brucellosis and Vaccine Safety. Infect Immun 2019;87. PMC6529653

14. Lacey CA, Mitchell WJ, Dadelahi AS, Skyberg JA. Caspase-1 and Caspase-11 Mediate Pyroptosis, Inflammation, and Control of Brucella Joint Infection. Infect Immun 2018;86. PMC6105886

15. Ruutu M, Thomas G, Steck R, Degli-Esposti MA, Zinkernagel MS, Alexander K, Velasco J, Strutton G, Tran A, Benham H, Rehaume L, Wilson RJ, Kikly K, Davies J, Pettit AR, Brown MA, McGuckin MA, Thomas R. beta-glucan triggers spondylarthritis and Crohn’s disease-like ileitis in SKG mice. Arthritis Rheum 2012;64:2211–22.

16. Benham H, Rehaume LM, Hasnain SZ, Velasco J, Baillet AC, Ruutu M, Kikly K, Wang R, Tseng HW, Thomas GP, Brown MA, Strutton G, McGuckin MA, Thomas R. Interleukin-23 mediates the intestinal response to microbial beta-1,3-glucan and the development of spondyloarthritis pathology in SKG mice. Arthritis Rheumatol 2014;66:1755–67.

17. Rehaume LM, Mondot S, Aguirre de Carcer D, Velasco J, Benham H, Hasnain SZ, Bowman J, Ruutu M, Hansbro PM, McGuckin MA, Morrison M, Thomas R. ZAP-70 genotype disrupts the relationship between microbiota and host, leading to spondyloarthritis and ileitis in SKG mice. Arthritis Rheumatol 2014;66:2780–92.

18. Baillet AC, Rehaume LM, Benham H, O’Meara CP, Armitage CW, Ruscher R, Brizard G, Harvie MC, Velasco J, Hansbro PM, Forrester JV, Degli-Esposti MA, Beagley KW, Thomas R. High Chlamydia Burden Promotes Tumor Necrosis Factor-Dependent Reactive Arthritis in SKG Mice. Arthritis Rheumatol 2015;67:1535–47.

19. Grillo MJ, Blasco JM, Gorvel JP, Moriyon I, Moreno E. What have we learned from brucellosis in the mouse model? Vet Res 2012;43:29. PMC3410789

20. Sakaguchi N, Takahashi T, Hata H, Nomura T, Tagami T, Yamazaki S, Sakihama T, Matsutani T, Negishi I, Nakatsuru S, Sakaguchi S. Altered thymic T-cell selection due to a mutation of the ZAP-70 gene causes autoimmune arthritis in mice. Nature 2003;426:454–60.

21. den Hartigh AB, Sun YH, Sondervan D, Heuvelmans N, Reinders MO, Ficht TA, Tsolis RM. Differential requirements for VirB1 and VirB2 during Brucella abortus infection. Infect Immun 2004;72:5143–9. PMC517456

22. Radhakrishnan GK, Yu Q, Harms JS, Splitter GA. Brucella TIR Domain-containing Protein Mimics Properties of the Toll-like Receptor Adaptor Protein TIRAP. J Biol Chem 2009;284:9892–8. 2665112

23. Chen CY, Nace GW, Irwin PL. A 6 x 6 drop plate method for simultaneous colony counting and MPN enumeration of Campylobacter jejuni, Listeria monocytogenes, and Escherichia coli. J Microbiol Methods 2003;55:475–9.

24. Sun Y, Connor MG, Pennington JM, Lawrenz MB. Development of bioluminescent bioreporters for in vitro and in vivo tracking of Yersinia pestis. PLoS One 2012;7:e47123. PMC3469486

25. Hayer S, Vervoordeldonk MJ, Denis MC, Armaka M, Hoffmann M, Bäcklund J, Nandakumar KS, Niederreiter B, Geka C, Fischer A, Woodworth N, Blüml S, Kollias G, Holmdahl R, Apparailly F, Koenders MI. ’SMASH’ recommendations for standardised microscopic arthritis scoring of histological sections from inflammatory arthritis animal models. Ann Rheum Dis 2021;80:714–26. PMC8142455

